# *β*-Cells as a Cell Factory for On-Demand Recombinant Protein Dosing: Harnessing the Neuroendocrine Cell Secretory Pathway for Controlled Release

**DOI:** 10.1101/2024.01.20.576492

**Authors:** Ella A. Thomson, Haixia Xu, Sooyeon Lee, Rayhan A. Lal, Justin P. Annes, Ada S. Y. Poon

## Abstract

This study explores the potential of utilizing *β*-cells, exemplified with R7T1 *β*-cell pseudoislets, as a transplantable cell factory for on-demand recombinant protein therapeutic delivery. While mammalian cell lines are widely used for *in vitro* protein production, the commonly utilized constitutive secretion pathway poses challenges to *in vivo* cell therapy, especially for delivering proteins requiring precise exposure kinetics. The proposed approach capitalizes on unique aspects of *β*-cells, including substantial vesicular protein storage capacity and electrochemically-regulated protein release, to facilitate timely and titratable *in vivo* therapeutic delivery. Examining a variety of strategies to acheive *β*-cell glucagon or glucagon-like peptide 1 (GLP-1) storage and secretion, we devised a flexible *β*-cell-based expression platform for efficient cellular peptide production and on-demand release. This platform utilizes the preproinsulin coding sequence as a template, wherein therapeutic peptides of interest (glucagon or GLP-1) are substituted for C-peptide while the A- and B-peptide insulin chains are mutated to prevent bio-active insulin production. This approach overcomes the challenge of efficient bio-active peptide expression by leveraging the endogenous *β*-cell peptide expression, translation, processing, storage and secretion machinery. Furthermore, *β*-cells provide a mechanism for scalable electyrochemnically-triggered peptide delivery. This transformative strategy, which may be extended to other proteins and peptide expression cassettes, holds significant promise for targeted and temporally controlled *in vivo* production and release of recombinant protein therapeutics. The study suggests potential applications in addressing challenges in metabolic disorders, blood disorders, and oncology. Future refinements may focus on optimizing vector design, peptide production, and *in vivo* adaptation.

## INTRODUCTION

Recombinantly produced proteins constitute a substantial portion of newly developed and approved drugs, and therapeutics (Sanchez-Garcia et al., 2016). While bacteria have historically served as cell factories for drug production, there has been a notable shift in recent years towards utilizing mammalian cell lines, including HEK293 cells, HeLa cells, plasma cells, and Chinese Hamster Ovary (CHO) cells (Tihanyi and Nyitray, 2020; Sharker and Rahman, 2021; Bryan et al., 2021; Arena et al., 2018). Despite this transition, therapeutic production primarily occurs in *in vitro* environments using large bioreactors. The transplantation of encapsulated cells is emerging as a promising avenue for durable treatment *via* implantable cell factories that provide a self-sustained, on-demand production and release of diverse therapeutics. This approach can be particularly crucial for recombinantly produced proteins characterized by a short half-life or rapid degradation, exemplified by peptides like glucagon, GLP-1, and parathyroid hormone. However, prevalent mammalian cell lines used for recombinant protein production lack granular protein storage and a secretory protein release pathway (Tihanyi and Nyitray, 2020; Sharker and Rahman, 2021). As a result, these cell lines exhibit a continuous protein secretion from cells, known as constitutive release (Burgess and Kelly, 1987), and lack the machinery necessary to release a substantial amount of protein within a short time frame, ranging from sub-minute to minute intervals. This presents challenges, particularly for proteins susceptible to rapid degradation, like glucagon and GLP-1, or those necessitating the release of a substantial protein quantity within a precise time frame, like glucagon for hypoglycemic recovery. Furthermore, therapeutic peptides are inefficiently translated and require post-translational modifications not performed via the constituitive secretion pathway, such as propeptide cleavage or C-terminal amidation, for maximal activity and stability (Chen et al., 2018; Germanos et al., 2021).

Here, we propose harnessing the potential of beta cells, specifically R7T1 *β*-cell pseudoislets, as an on-demand cell factory for recombinant proteins. Beta cells stand out as an ideal target cell line due to their possession of peptide processing machinery, vesicle protein storage and a regulated vescicular fusion (cargo release). The proteins, once translated and processed, are stored within vesicles within the cells (Burgess and Kelly, 1987) (Fig. 1). The process of membrane depolarization, induced by stimuli such as glucose, KCl (Fig. 1), or electrical signals, propels the exocytosis of the stored protein from the cell. This intricate mechanism facilitates the prompt and controlled release of a substantial quantity of stored protein, responding swiftly to demand within a sub-minute to minute time frame. Furthermore, R7T1 *β* cells exhibit reversible immortalization and the capability to organize into pseudoislet clusters, thereby providing a mechanism for manipulating cell numbers and cell density within transplanted encapsulation. Furthermore, electrical coupling of R7T1 *β*-cell pseudoislets provides a mechanism for synchronized protein release. In this work, we demonstrate the use of *β*-cells for the synthesis of glucagon and GLP-1, highlighting their controlled release along the secretory release pathway of *β*-cells.

**Figure 1.**
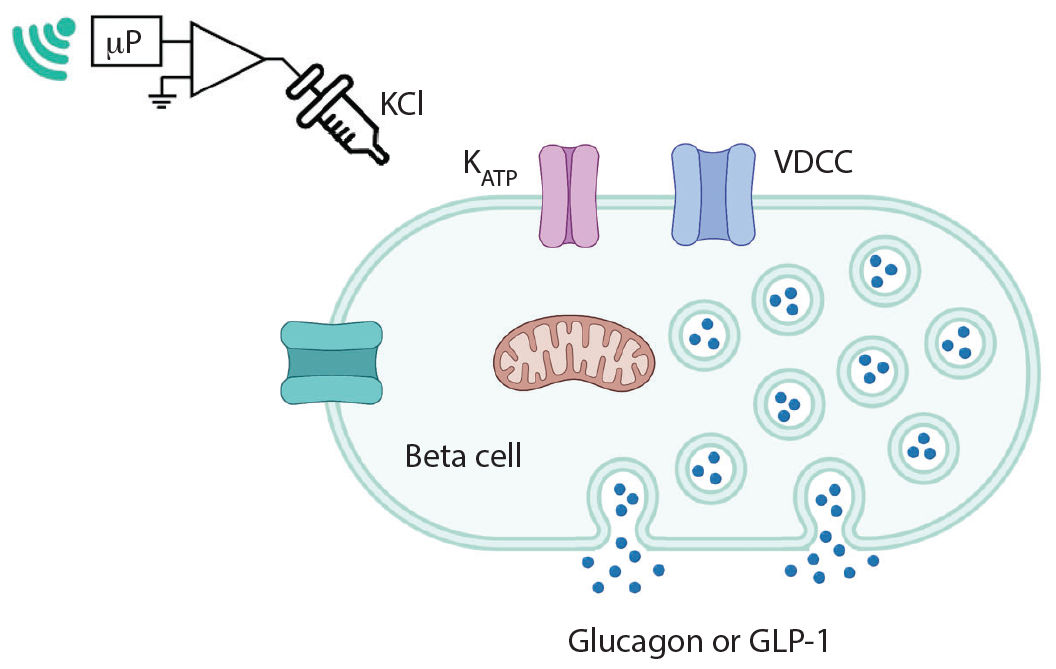
Illustration of vesicle protein storage of a beta cell and the activation of its secretory protein release pathway using KCl stimulation.

Beta cells natively produce and process insulin. Insulin is initially translated as preproinsulin, comprising a signal peptide (SP), A-chain, B-chain and C-peptide (Fig. 2). The signal peptide guides preproinsulin into the endoplasmic reticulum for processing. The interaction of the signal peptide with cytosolic ribonucleoprotein signal recognition particles enables translocation into the lumen across the rough endoplasmic reticulum (Fu et al., 2013). Subsequently, the signal peptide is cleaved by signal peptidase, resulting in the formation of proinsulin (Fig. 2). Proinsulin undergoes folding, forming three disulfide bonds, and is then transported to the Golgi apparatus. Within the Golgi apparatus, it enters secretory vesicles, where enzymes cleave proinsulin into mature insulin and C-peptide (Fig. 2). Both mature insulin, composed of the A-chain and B-chain linked by disulfide bonds, and C-peptide are stored in vesicles within the cell.

**Figure 2.**
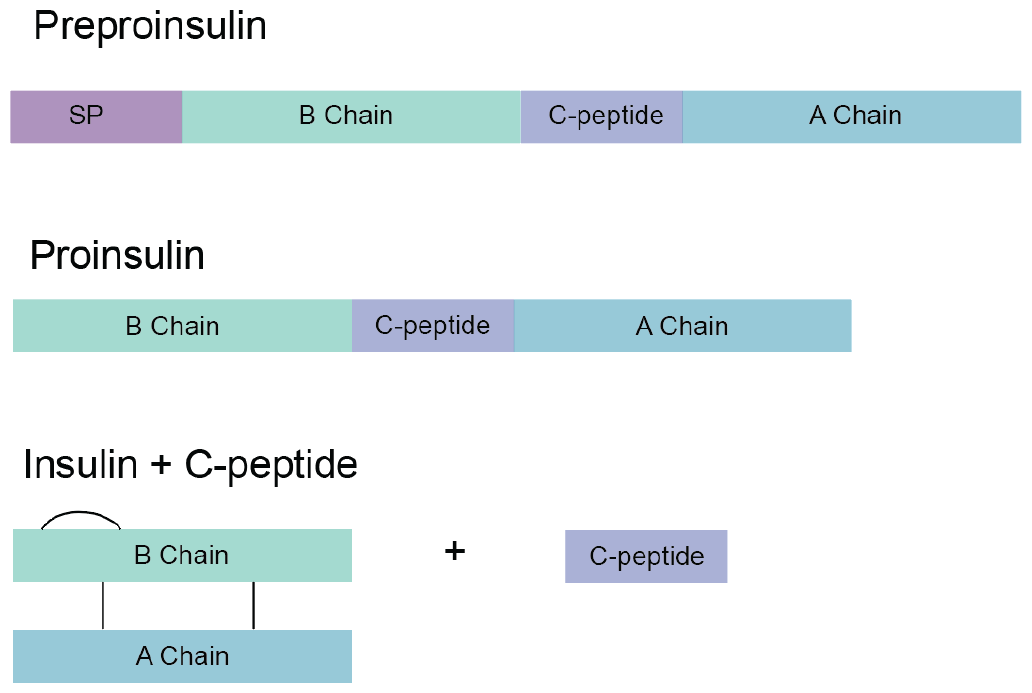
A visual representation of process of insulin production and processing in beta cells.

To direct the release of the protein of interest by *β*-cells, we designed DNA vectors in which the C-peptide portion of proinsulin was substituted with our protein of interest. Additionally, mutations were introduced into the A-chain and B-chain of insulin to avoid co-secretion of bio-active insulin with the protein of interest. Additionally, we used CRISPR to disrupt genomically encoded insulin to prevent endogenous insulin production. Finally, using a luciferase-based expression platform, we validated that the engineered vectors follow the secretory pathway upon release.

## METHODS AND MATERIALS

### Culturing and characterization of R7T1 *β* cells and HEK293 cells

Immortalized mouse R7T1 *β*-cells were cultured in DMEM high glucose containing 2 mM L-glutamine and 1 mM sodium pyruvate, with 10% FBS, 1% penicillin/streptomycin, and 1 *μ*g/mL doxycycline (Sigma-Aldrich, St. Louis, MO, USA). R7T1 *β* cells (passage number 10–34) were growth arrested for 5 days with doxycycline-free medium supplemented with tetracycline free fetal bovine serum (Gibco A4736301). To form pseudoislets, growth arrested R7T1 *β* cells were plated at 300,000 cells per well in a 12-well non-adherent plate and after 16–24 hours, pseudoislets were picked and counted for experiments. R7T1 *β* cells were characterized by comparing growth arrested and proliferating insulin secretion.

Cells were growth arrested for 5 days by removing doxycycline from culture media. Proliferating cells were not growth arrested and were culture in media supplemented with doxycycline for a five-day period. 300,000 cells per well were plated in a 12-well cell culture plate and cultured in DMEM low glucose (with or without doxycycline) for 24 hours. Media was then replaced with high glucose media and supernatant samples were collected after 30 minutes. Insulin content in supernatant was evaluated by rodent insulin ELISA (ALPCO STELLUX 80-INSMR-CH01).

HEK293 cells were cultured in DMEM high glucose containing 2 mM L-glutamine and 1 mM sodium pyruvate, with 10% FBS and 1% penicillin/streptomycin.

### CRISPR Cas9 insulin knockout

CRISPR Cas9 (Jiang et al., 2017; Hsu et al., 2014) was used to disrupt the Ins1 and Ins2 genes of R7T1 *β* cells. Initially, a stable R7T1 *β* cell line expressing Cas9 was generated through lentiviral transduction, followed by the selection of cells resistant to blasticidin. Two guides were designed to target the Ins1 and Ins2 genes, with assessment of potential off-target effects using CHOPCHOP. The guide sequences are detailed in Table 1. These guides were then mapped onto genomic DNA, and primers were designed for subsequent TIDE analysis. Genomic DNA extracted from the target cells was aligned with the sequenced genome. Figure 3 illustrates the precise locations of the two guides on the Ins1 gene. Oligos for each guide were synthesized (IDT).

**Table 1.**
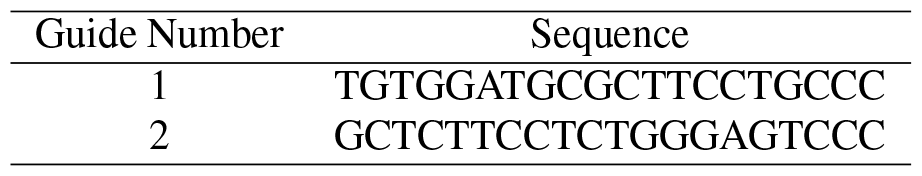
Guide sequences used to knockout Ins1 and Ins2 genes in R7T1 beta cells.

**Figure 3.**
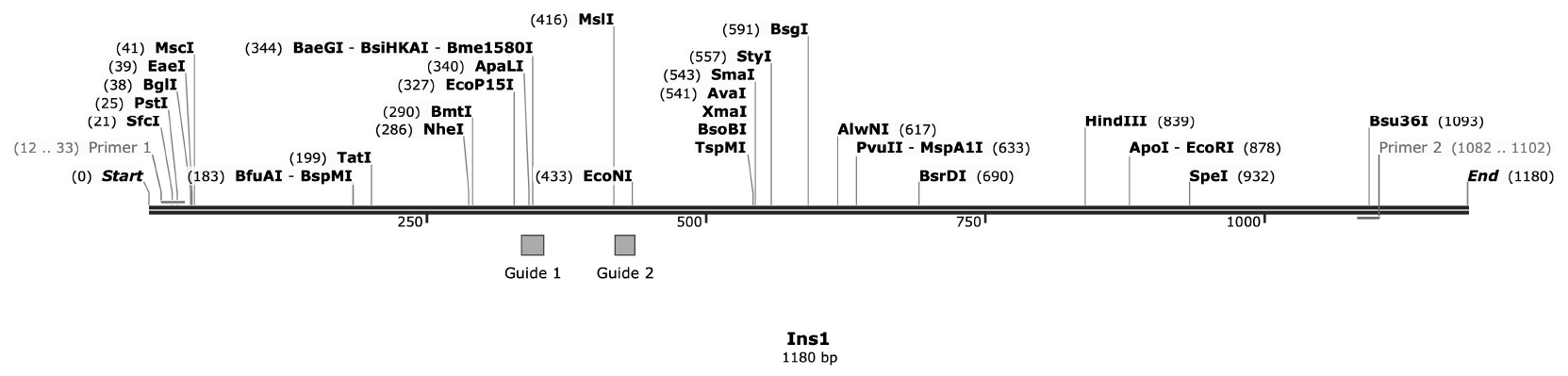
Locations of guides in Ins1 gene.

The guides were integrated into a lentiviral guide expression plasmid (pcmb320, a gift from Michael Bassik (Addgene plasmid 89359). The knockout process was executed via lentiviral transduction. Viral packaging of each guide was performed using HEK293 cells transfected with polyethylenimine (PEI) diluted in reduced serum OptiMEM. Subsequently, the Cas9-expressing R7T1 *β* cells were sequentially exposed to the conditioned media of transfected HEK293 cell media for 24 hours (viral conditioned media collected at 48 and 72 hours). Seventy-two hours post-transduction, puromycin selection was initiated. One week after initiating puromycin selection, flow cytometry (FACS) sorting was performed to select for mCherry expression. Following sorting, cells were maintained under puromycin and blasticidin selection. Immunostaining and ELISA were conducted to assess intracellular insulin content and confirm disrupted insulin expression.

### Immunofluorescent staining and quantification of insulin expression in fixed cells

Cells underwent fixation using a solution of 4% paraformaldehyde in phosphate-buffered saline. Staining was carried out using an anti-insulin primary antibody at a 1:500 dilution. Hoechst was employed as both a nuclear stain and for quantifying cell number. Subsequently, the fixed and stained cells were preserved in PBS at 4 *°*C. The quantification of insulin intensity was conducted using Cellomics Software (ArrayScan).

### Insulin quantification in R7T1 *β* cells

The R7T1 *β* cells were seeded in a 12-well cell culture dish. After 24 hours, the media was aspirated, and the cells were washed with PBS. Subsequently, the cells were lysed in 50-*μ*l RIPA buffer (50 mM tris HCl, 150 mM NaCl, 1% NP-40, 0.1% SDS, 0.5% Deoxycholic acid). The plates were frozen at *−* 20 *°*C for a minimum of 16 hours. After thawing, the cells were scraped from the bottom of the cell culture plate.

The resulting samples were collected in Eppendorf tubes and centrifuged at 14,000 RPM for 20 minutes. The supernatants were then collected and subjected to dilution. The insulin content of the samples was assessed using ELISA (ALPCO Stellux).

### Design and optimization of glucagon and GLP-1 expression vectors

The design and production of Glucagon and GLP-1 expression vectors were conducted using SnapGene, and the vectors were synthesized by Twist. A bacterial transformation was carried out using competent Stbl3 E. coli, followed by a midiprep procedure. Subsequently, DNA sequencing was performed at MCLAB.

In the initial phase, nine glucagon vector designs were created, encompassing variations in promoters, vector backbones, and selection markers. Promoters included CMV and SFFV, while backbones consisted of non-viral and 3rd generation lentiviral vectors. Selection markers incorporated RFP, GFP, and hygromycin resistance, added through either a 2A linker or IRES. Additional modifications involved the inclusion of the WPRE sequence, Kozak sequence, and the use of both the native glucagon signal peptide and an optimized signal peptide sequence. Table 2 summaries the vector designs. Additionally, the gateway cloning protocol was employed to transfer the DNA insert into a different backbone.

**Table 2.** Summary of various versions of glucagon vector design.

| Vector | Promoter | Backbone | Glucagon Variant | Linker | Tag |
| --- | --- | --- | --- | --- | --- |
| 1 | SFFV | Lentiviral | Native | IRES | GFP |
| 2 | SFFV | Lentiviral | Native | T2A | RFP |
| 3 | SFFV | Lentiviral | Native + UTR | IRES | GFP |
| 4 | CMV | Non-viral | Native + Glucagon SP | None | None |
| 5 | CMV | Non-viral | Native + Optimized SP | None | None |
| 6 | CMV | Non-viral | Glicentin | None | None |
| 7 | CMV | Non-viral | Native | None | None |
| 8 | SFFV | Lentiviral | Native | None | None |
| 9 | CMV | Lentiviral | Native | None | Hygromycin |

The ultimate vector design featured a CMV promoter and a modified proinsulin sequence. The altered proinsulin sequence encompassed the preproinsulin signal peptide, A-chain, and B-chain. Notably, the C-peptide sequence was replaced with the glucagon sequence. This DNA insert was employed in two distinct backbones–a mammalian expression vector and a 3rd generation lentiviral vector for stable line development. The latter included an antibiotic resistance selection marker.

### Transfection of R7T1 *β* cells and HEK293 cells

Twenty-four hours prior to transfection, cells were seeded either in a 15-cm dish or a 12-well adherent cell culture plate. Transfection with Lipofectamine 2000 occurred at approximately 80% confluence for R7T1 *β* cells and 50% confluence for HEK293 cells, utilizing Opti-MEM reduced serum media as per the manufacturer’s specifications. The cells were co-transfected with pmaxGFP and either glucagon, GLP-1, or dual-luciferase (proinsulin renilla or CMV firefly) expression vectors. Control samples received pmaxGFP (with equivalent total DNA used for all transfections). A media change was conducted 4-6 hours post-transfection.

### Experimental configuration for stimulated protein secretion in transfected R7T1 *β* cells and HEK293 cells

Secretion experiments were conducted 48 hours post-transfection. R7T1 *β* cells underwent a 24-hour maintenance period in DMEM low glucose containing 2 mM L-glutamine and 1 mM sodium pyruvate, with 10% FBS, 1% penicillin/streptomycin, and 1 *μ*g/mL doxycycline (Sigma-Aldrich, St. Louis, MO, USA). Meanwhile, HEK293 cells were sustained in DMEM low glucose containing 2 mM L-glutamine and 1 mM sodium pyruvate, with 10% FBS and 1% penicillin/streptomycin. Just before the secretion experiment, each well underwent three wash steps in 1 ml of DMEM low glucose containing 2 mM L-glutamine and 1 mM sodium pyruvate, with 10% FBS and 1% penicillin/streptomycin.

The basal media for the experiment was DMEM low glucose. The stimulation media was additionally enriched with 40 mM KCl. A 0.4-ml media volume was employed for the experiments. After 15 minutes in the fresh media, a 0.2-ml conditioned media sample was collected for each well. These samples underwent centrifugation for 2 minutes at 2,000 RPM, and a 0.15-ml supernatant sample was collected. Samples for GLP-1 and glucagon release were immediately measured by ELISA (Crystal Chem and ABCAM). Conversely, samples for human insulin measurements were stored at *−* 20 *°*C and subsequently assessed by ELISA after undergoing one thaw cycle (ALPCO Stellux).

### Dual Luciferase secretion experiments

Secretion experiments were conducted 48 hours post-transfection. R7T1 *β* cells were pre-incubated in DMEM low glucose media for 24 hours before the experiments, while HEK293 cells were maintained in standard cell culture media until the experiment. Immediately prior to the experiment, each well underwent three wash steps with 1 ml of DMEM low glucose media. The basal media for the experiment was DMEM low glucose media, and the stimulation media was additionally supplemented with 40 mM KCl. A media volume of 0.4 ml was used for the experiments. After 15 minutes in the fresh media, a 0.2-ml conditioned media sample was collected for each well. These samples underwent centrifugation for 2 minutes at 2,000 RPM, and a 0.15-ml supernatant sample was collected. Following this, all wells were rinsed with Phosphate Buffered Saline, and cells were lysed in 0.5 ml of Luciferase Lysis Buffer (25 mM Tris/phosphate, 4 mM EGTA, 1% Triton X-100, 10% glycerol, 2 mM dithiothreitol). Supernatant and lysate samples were stored at 20 *°*C and subsequently evaluated by ELISA after one thaw cycle (ALPCO Stellux). For luminescence evaluation, 0.02 ml of sample was assessed with 0.08 ml of Renilla luciferase buffer (1.1 M NaCl, 2.2 mM EDTA, 220 mM KxPO4, 0.44 mg/ml BSA, 1.33 mM NaN3, and 1.43 *μ*M ctz) and Firefly luciferase buffer (25 mM glycylglycine, 15 mM KxPO4, 4 mM EGTA, 2 mM ATP, 1 mM DTT, 15 mM MgSO4, 100 *μ*M CoA, and 75 *μ*M Luciferin) using a 0.5-second integration time (Kalwat et al., 2016).

### *in Vitro* Glucagon measurements with encapsulated Pseudoislets

Twenty-four hours post-transfection, growth arrest was induced in R7T1 cells, followed by the formation and selection of pseudoislets at 48 hours post-transfection. PET membranes were affixed to cell culture inserts using PET sealant (Hotmelt) and positioned in a 12-well cell culture plate containing 3 ml DMEM supplemented with 1 IU/mL Penicillin, 1 *μ*g/mL Streptomycin, and 1 mM sodium pyruvate to moisten the membrane. The insert was then filled with 300 *μ*l of pseudoislets suspended in supplemented DMEM. For KCl-stimulated cells, Potassium Chloride was introduced into the cell culture insert to achieve a final concentration of 40 mM. After 15 minutes, media samples were collected from inside the encapsulation to measure the total secreted insulin, while diffusion samples were collected from the media in the cell culture plate. In the case of insulin release induced by pressure, a piezoelectric pump and controller were attached to the cell culture insert and activated for 30 seconds with a pressure of 11 kPa. Following pump deactivation, a sample was collected from the media in the cell culture plate, and the volume of media outside the encapsulation was also measured. The concentration of glucagon was determined by ELISA (Crystal Chem), and the total released glucagon was subsequently calculated.

## RESULTS

### CRISPR-mediated insulin knockout in R7T1 *β*-cells

Utilizing CRISPR Cas9, we generated a line of insulin knockout R7T1-Cas9 *β*-cells to prevent cosecretion of insulin and target peptides in subsequent experiments. Following transduction of R7T1-Cas9 *β*-cells with INS1- and INS2-targeted guides, antibiotic selection (puromycin) and flow cytometry sorting (mCherry selection for transduced cells), we evaluated insulin expression levels in wild-type and knockout R7T1 *β* cells by immunostaining and ELISA. Immunostaining of knockout R7T1 *β* cells revealed a substantially reduced insulin-positive cell population compared to wild-type cells (Fig. 4A). Concordantly, knockout R7T1 *β* cells demonstrated substantially reduced insulin expression compared to wild-typ R7T1 cells as determined by ELISA (Fig. 4B). These data confirmed sucsesful impairment of insulin expression in the targeted R7T1-Cas9 *β*-cells.

**Figure 4.**
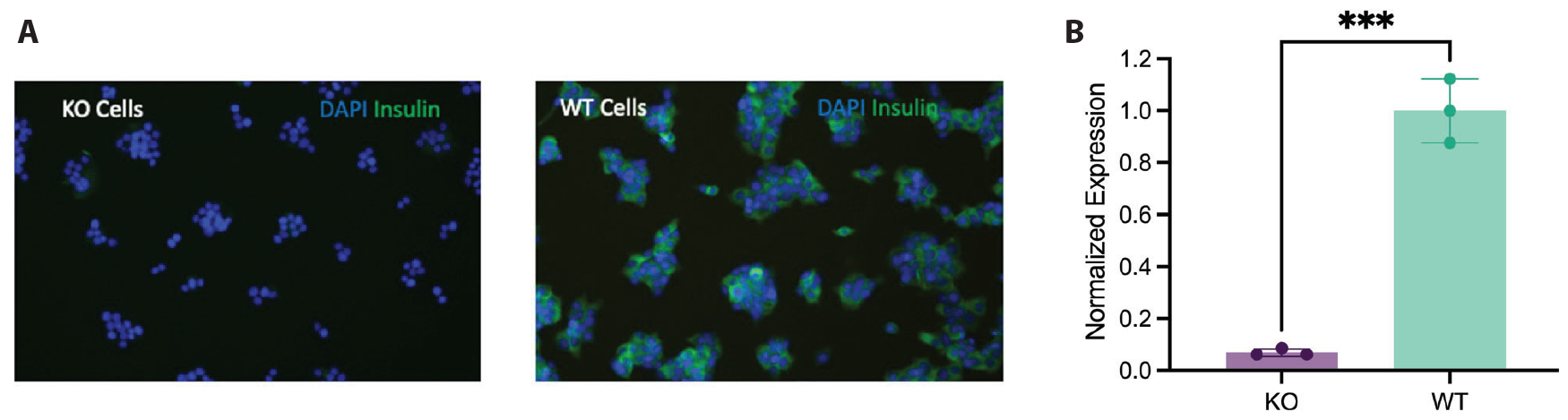
Insulin knockout in R7T1 *β* cells. CRISPR Cas9 knockout results in decreased expression of insulin in R7T1 *β* cells evaluated by (**A**) immunostaining and (**B**) intracellular lysate ELISA.

### Engineered proinsulin vector for glucagon secretion

To harness the insulin processing capabilities of beta cells (Fig. 2), we engineered the proinsulin vector to replace the C-peptide sequence with the glucagon sequence. In native preproinsulin, proinsulin undergoes processing to insulin through the enzymes prohormone convertase 2 (PC2) and prohormone convertase 1/3 (PC1/3) (Fig. 5A). Additional enzyme cleavage sites for PC2 are incorporated on both sides of the glucagon sequence to ensure the cleavage of additional amino acids, thereby eliminating any remaining linkers attached to the glucagon peptide (Fig. 5A). To prevent the co-secretion of active insulin and glucagon, two mutations were introduced to the A-chain and B-chain of insulin, rendering it a non-bioactive form. Consequently, the final vector was meticulously designed to co-secrete an inactive form of insulin alongside an active form of glucagon (Fig. 5A).

**Figure 5.**
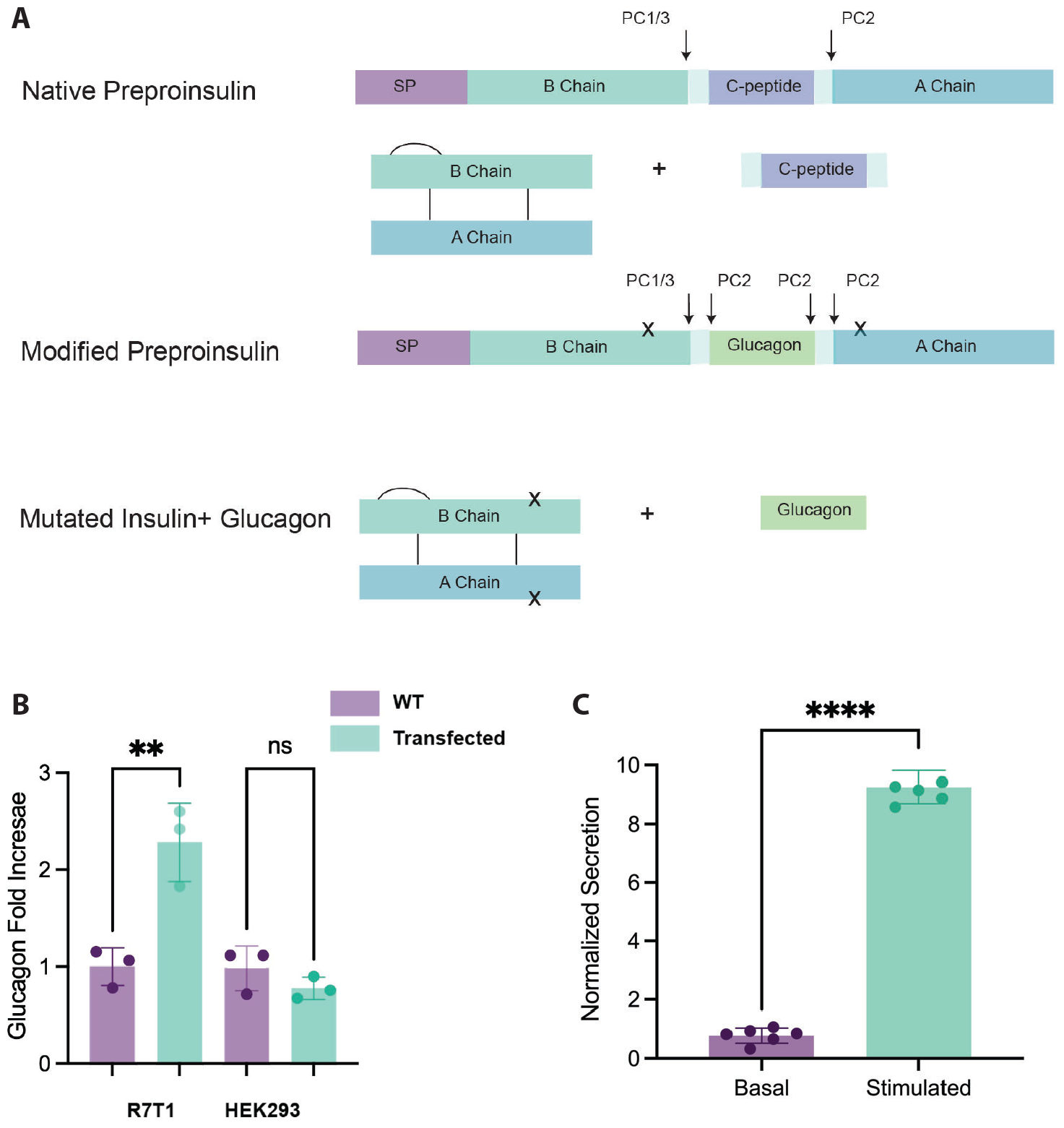
Engineering the proinsulin vector for glucagon secretion. (**A**) Design of the proinsulin glucagon vector targeted for R7T1 enzyme processing. (**B**) Engineered proinsulin glucagon vector increased secretion level of glucagon from transfected R7T1 cells but not from transfected HEK293 cells. (**C**) KCl stimulation increases glucagon secretion from transfected *β* cells.

Using the engineered proinsulin vector, we assessed the secretion of glucagon in transfected R7T1 *β* cells and compared to control cells. Transfected cells exhibited a heightened level of glucagon secretion compared to untransfected controls (Fig. 5B). Despite the introduction of insulin mutations resulting in decreased secreted insulin levels, it had no impact on the secretion levels of our protein of interest. Additionally, HEK293 were transfected with the same engineered proinsulin vector, and in comparison to control cells, no increased glucagon levels were detected in the conditioned media (Fig. 5B). This outcome suggests that the engineered proinsulin vector is undergoing processing and cleavage as intended, given that HEK293 cells lack the necessary enzymes (PC1/3 and PC2) to process proinsulin. Upon cell stimulation via KCl-driven membrane depolarization, secreted glucagon levels were nearly 8-fold higher than in unstimulated controls (Fig. 5C). This finding suggests that stimulation can effectively trigger the release of a bolus of glucagon from our genetically engineered cell line, indicating that glucagon is likely being released along the beta cell secretory pathway.

### Validation of constitutive vs. secretory release pathways

To validate the release pathways of the engineered proinsulin vectors, R7T1 *β* cells and HEK293 cells were co-transfected with two luciferase vectors: the proinsulin renilla and a CMV firefly (Fig. 6A). The signal peptide in the proinsulin renilla vector should direct the renilla luciferase along the secretory pathway, while the firefly vector lacks a signal peptide and should be directed to the constitutive pathway only. Additionally, the firefly expression level serves to normalize transfection efficiency between the various cell lines and conditions. Upon KCl stimulation, an increase in renilla luciferase secretion from R7T1 *β* cells is expected, but not from HEK293 cells (Fig. 6B)

**Figure 6.**
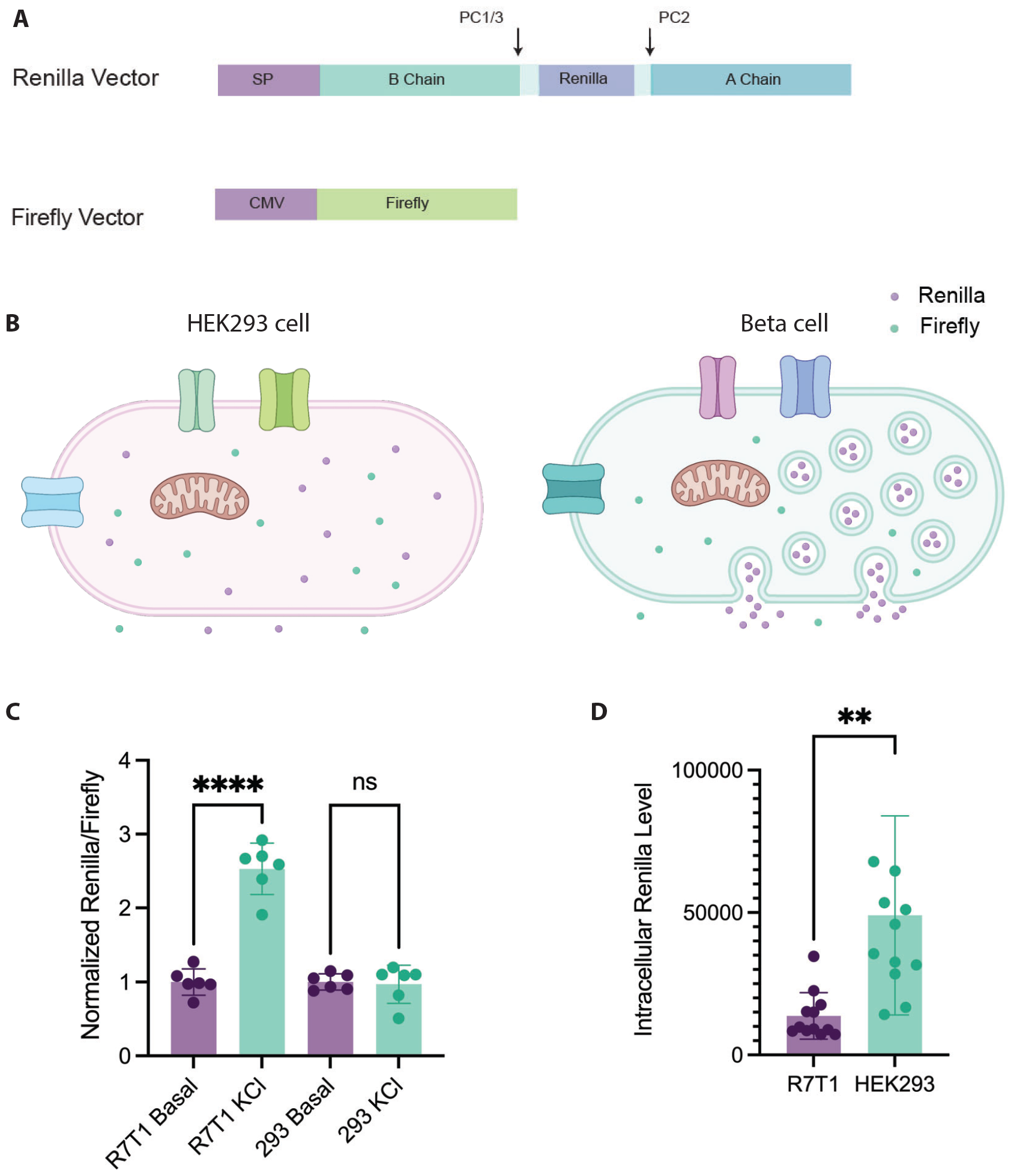
Validation of constitutive versus secretory release pathways. (**A**) A visual representation of the Firefly and renilla luciferase vector insert designs. (**B**) A visual illustration of our hypothesis on the secretion of Firefly and renilla luciferase upon KCl stimulation in transfected HEK293 cells and *β* cells. (**C**) Normalized secretion levels of renilla luciferase upon KCl stimulation in transfected HEK293 cells and R7T1 *β* cells. (**D**) Intracellular levels of renilla luciferase in transfected HEK293 cells and R7T1 *β* cells.

In R7T1 *β* cells, KCl stimulation resulted in an increased secretion of renilla luciferase, normalized for transfection efficiency (Fig. 6C). Conversely, in HEK293 cells, KCl stimlation did not result in an increase secretion of renilla luciferase relative to basal levels, normalized for transfection efficiency (Fig. 6C). This indicates that the proinsulin vector is directed to the secretory pathway in the R7T1 *β* cells but is only constitutively released in the HEK293 cells. Importantly, the intracellular level of renilla luciferase was higher in HEK293 cells than in R7T1 *β* cells (Fig. 6D). This indicates that the difference observed in the previous experiment (Fig. 5B) is not due to the inability of HEK293 cells to translate the engineered proinsulin vector but rather due to the lack of granular protein storage in HEK293 cells.

### Utilizing the engineered proinsulin vector as a platform for multiple peptide releases

Beyond glucagon, we modified the proinsulin vector by substituting the C-peptide sequence with the GLP-1 sequence. Similar to the glucagon vector, we observed increased secretion of GLP-1 in transfected cells upon KCl stimulation (Fig. 7). This suggests that the engineered proinsulin vector is versatile and can be employed for the production of various peptides with stimulated secretion.

**Figure 7.**
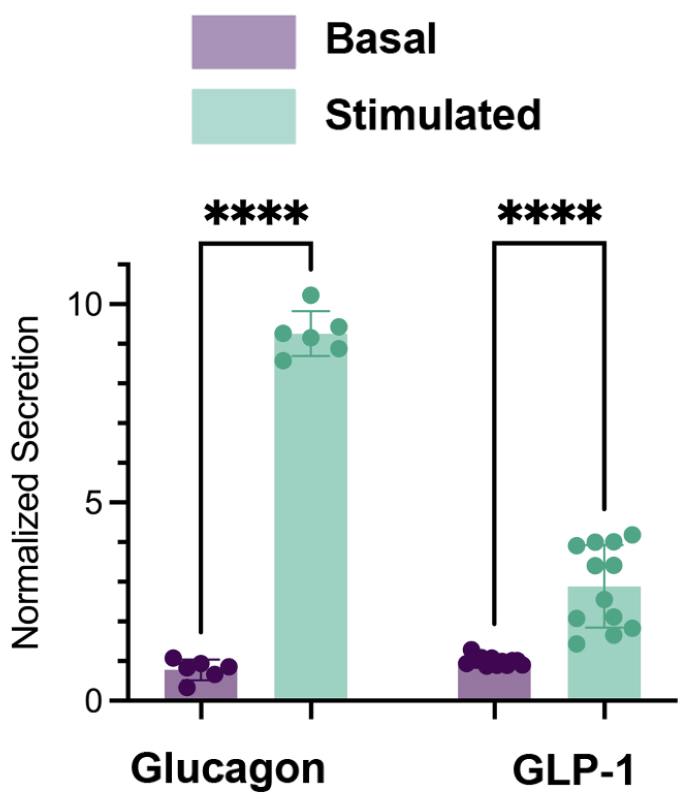
KCl stimulation results in elevated secretion of glucagon and GLP-1 from R7T1 cells transfected with the engineered proinsulin vectors.

## DISCUSSIONS

In this study, we utilized the insulin processing capabilities of beta cells to demonstrate their potential for producing and secreting glucagon and GLP-1. By engineering a proinsulin vector in which the C-peptide section is substituted with glucagon or GLP-1, we observed increased secretion of the targeted proteins in response to KCl stimulation. This demonstrates the potential to synthetically produce, package, and store peptides in R7T1 *β* cells for on-demand secretion. Utilizing a luciferase-based expression platform, we verified that the engineered vectors followed the native secretory granule pathway used for insulin biosynthesis and release. This methodology presents an opportunity for further refinement to enhance peptide delivery efficacy. Future investigations could focus on optimizing vector design by improving promoter strength and incorporating additional beta-cell processing enzymes, such as carboxypeptidases and the C-terminal amidating activity of Peptidyl-glycine alpha-amidating monooxygenase (PAM), to enhance peptide stability and bioactivity. Additionally, exploring the introduction of multiple copies of relevant genes could maximize peptide production levels and enable the simultaneous production of multiple distinct peptide therapeutics.

The implications of this work extend beyond the immediate scope, potentially offering a transformative approach to the production and delivery of recombinantly produced proteins. Beta cells, with their vesicle peptide storage and the ability to release stored peptides in response to membrane depolarization, provide an advantageous platform. The capacity of R7T1 *β* cells to form pseudoislets further enhances the maximum achievable encapsulated cell density for an implantable device. These unique features make it possible to develop an *in vivo* cell factory for recombinantly produced proteins. The presented approach enables targeted and temporally controlled production and release of recombinantly produced therapeutics, with broad applications in metabolic disorders, blood disorders, and oncology. Notably, it opens a pathway for the production and temporally controlled delivery of therapeutic recombinant peptides.

## ACKNOWLEDGMENTS

We are thankful to the funding support from Stanford Bio-X Interdisciplinary Initiatives Seed Grants Program (IIP) (R10-56), Stanford Graduate Fellowship, and Stanford Bio-X Bowes Fellowship.

## Notes

### Competing Interest Statement

E. A. T., R. A. L.,, J. P. A., and A.S.Y.P. are co-inventors of a patent covering the work described in the manuscript filed by Stanford.

